# Formulation, Characterization, and *in vivo* Immunogenicity of Heat-Stabilized Dissolvable Microneedles Containing a Novel VLP Vaccine

**DOI:** 10.1101/2024.12.16.628763

**Authors:** Aidan Leyba, Alexandra Francian, Mohammad Razjmoo, Amelia Bierle, Ranjith Janardhana, Nathan Jackson, Bryce Chackerian, Pavan Muttil

**Author notes:** AL and AF contributed equally to this paper.

## Abstract

Since its introduction, vaccination has heavily improved health outcomes. However, implementing vaccination efforts can be challenging, particularly in low and middle-income countries with warmer climates. Microneedle technology has been developed for its simple and relatively painless applications of vaccines. However, no microneedle vaccine has yet been approved by the FDA. A few hurdles must be overcome, including the need to evaluate the safety and biocompatibility of the polymer used to fabricate these microneedles. Additionally, it is important to demonstrate reliable immune responses comparable to or better than those achieved through traditional administration routes. Scalability in manufacturing and the ability to maintain vaccine potency during storage and transportation are also critical factors. In this study, we developed vaccine-loaded dissolvable microneedles that showed preclinical immunogenicity after storage in extreme conditions. We developed our microneedles using the conventional micromolding technique with polyacrylic acid (PAA) polymer, incorporating a novel virus-like particle (VLP) vaccine targeting arboviruses. We performed characterization studies on these microneedles to assess needle sharpness, skin insertion force, and VLP integrity. We also investigated the thermostability of the vaccine after storing the microneedles at elevated temperatures for approximately 140 days. Finally, we evaluated the immunogenicity of this vaccine in mice, comparing transdermal (microneedle) with intramuscular (hypodermic needle) administration. We successfully fabricated and characterized VLP-loaded microneedles that could penetrate the skin and maintain vaccine integrity even after exposure to extreme storage conditions. These microneedles also elicited robust and long-lasting antibody responses similar to those achieved with intramuscular administration.

## Introduction

Vaccination has become a cornerstone of modern medicine over the past 200 years, enabling us to combat numerous infectious diseases and save millions of lives^1,2^. While immunization efforts have been successful throughout history, the process of immunization, encompassing storage, transportation, distribution, and administration of vaccines, has grown increasingly complex with the introduction of new vaccines. Most vaccines require continuous cold chain management during storage and transportation to maintain vaccine potency; this necessitates adequately trained staff and specialized equipment to keep vaccines in temperature-controlled environments from the moment they leave the manufacturing site until they are administered to individuals worldwide^3,4,5^. For instance, during the peak of the COVID-19 pandemic, low- and middle-income countries (LMICs) faced significant challenges in storing and distributing mRNA vaccines, making it difficult to vaccinate their populations^6^.

Most vaccines are administered using hypodermic needles, hindering global vaccination efforts. Proper administration requires trained healthcare workers; a shortage of these professionals, observed in LMICs, can lead to accidental needlestick injuries and the transmission of infections^7^. Such injuries are often caused by not following standard vaccination protocols, such as recapping needles after use^8^. Further, needle phobia affects many individuals; in the US alone, over 11 million people have a fear of needles^9^, which results in decreased compliance and inadequate vaccination coverage. LMICs will continue to face obstacles in achieving widespread immunization if current vaccination strategies remain unchanged. For instance, population growth and poor transportation networks will limit the timely delivery of vaccines in remote areas, forcing healthcare workers to travel long distances under challenging conditions, such as extreme temperatures and humidity, potentially compromising vaccine potency^10^.

Various strategies have been explored to address the limitations of needle vaccination, including the evaluation of oral, transdermal, and inhaled routes of administration^11^. Microneedles (MNs) offer a minimally invasive immunization method that delivers vaccines through the skin (transdermal delivery). MNs consist of arrays of micron-sized needles, providing a pain-free vaccine delivery option. Additionally, they present several advantages over traditional vaccination routes (such as intramuscular and subcutaneous injections), including reducing the fear associated with needle use and eliminating concerns about needlestick injuries. MN administration is straightforward and can be self-administered, which removes the need for trained personnel for administration^12^. Unlike liquid-based vaccines, those incorporated in MNs can be optimized to minimize cold-chain requirements due to their dry state and the materials used in manufacturing (such as sugars and polymers)^13^. Mistilis et al. showed that the influenza vaccine remains effective within MNs; the hemagglutinin activity was retained within MNs after storage at room temperature for 6 months^14^.

Different types of MNs have been used for drug and vaccine delivery^15^. Solid MNs create micro-pores in the skin, allowing the cargo to be delivered beneath the skin. Vaccines can also be coated onto the MN surface. Hollow MNs release their contents through tiny pores located at the needle tips. Hydrogel-based MNs swell when inserted into the skin, enabling controlled release of the contents^16^. Lastly, dissolvable MNs consist of safe and biocompatible polymers that encapsulate the cargo and dissolve under the skin after application^17^. The design and fabrication of dissolvable MNs and their simple application process make them highly desirable, especially since they do not create sharp waste after use^18^. From a regulatory perspective, dissolvable MNs are classified as medical devices and combination products (vaccine and polymer). Regulatory considerations for MNs include needle length, sharpness, ease of penetration into the skin, and whether they are administered manually or with an applicator^19^.

Our study evaluated the preclinical immunogenicity of thermostable and dissolvable MNs containing a novel vaccine. The micron-sized needles penetrate the skin and dissolve beneath its surface in the interstitial fluid, exposing the vaccine to antigen-presenting cells (APCs). APCs, such as dendritic cells (DCs) and Langerhans cells (LCs), are present in the superficial layers of the skin (epidermis and dermis), where the needles dissolve to release the vaccine^20^. The site of delivery potentially allows the vaccine to reach the nearest draining lymph nodes and elicit an immune response through lymphocyte stimulation and antibody production. Zaric et al. demonstrated that LCs are critical immune cells that elicit an immune response when vaccines are delivered using MNs^21^.

In this proof-of-concept study, we loaded our MNs with a novel Qβ bacteriophage virus-like particle (VLP) vaccine targeting sialokinin (SLK), a salivary peptide present in the *Aedes aegypti* mosquito that promotes infection by mosquito-borne viruses. VLPs are highly multivalent structures that serve effectively as vaccine platforms, eliciting strong antibody responses against various antigens. We examined the force required by the MNs to penetrate the skin barrier and their dissolving capabilities beneath the mouse skin. Finally, we assessed our VLP-MNs to demonstrate the thermostability of the vaccine and the MN vaccine delivery platform in producing a robust and long-lasting immune response in mice.

## Methods

### Materials

Dissolvable MNs were made from 35% poly(acrylic acid) (PAA; average MW ∼250,000) polymer using a micro-molding technique. Polydimethylsiloxane (PDMS) microneedle molds (10X10 array; height-600µm; Base-200µm) were used to make the MNs and were purchased from MicroPoint Technologies Pte Ltd (Singapore). Each PDMS mold contained an array of 10×10 wells that produced 100 needles.

### VLP Preparation

Qβ bacteriophage VLPs were produced in *Escherichia coli* using previously described methods^22^. Prior to peptide modification, WT Qβ VLPs were depleted of endotoxin by 4 rounds of phase separation using Triton X-114 (Sigma-Aldrich), as previously described^23^. The SLK peptide was synthesized *de novo* (GenScript) and modified to contain a C-terminal linker sequence *gly-gly-gly-cys* (NTGDKFYGLM-GGGC). SLK was conjugated to the surface lysine residues on Qβ VLPs using the bidirectional crosslinker, succinyl 6-[(β-maleimidopropionamido)hexanoate], or SMPH (ThermoFisher Scientific). VLP-peptide conjugation efficiency was assessed by SDS-PAGE, where a shift in molecular weight was observed by the addition of the peptide.

### MN Preparation

We formulated the VLP-containing MNs through a three-stage process that was completed within 24 hours (**Figure 1**). In the first stage, we prepared a 29% stock solution of PAA, which was gently vortex-mixed with VLPs (conjugated to SLK) to form a homogenous suspension. The VLP concentration used was 1.0mg/mL in phosphate-buffered saline (PBS). To achieve a vaccine dose of 10 µg per MN, we added 10 µL of VLPs to 30 µL of the PAA stock solution (**Figure 1A**). The second stage involved carefully pipetting 40 µL of this VLP-PAA suspension (comprising the needle layer) into PDMS molds, followed by centrifugation (Eppendorf Centrifuge 5810 R 15-amp version) (60 minutes at 30°C; 1350 RCF) (**Figure 1B**). The centrifugation allowed the polymeric VLP suspension to fill the needle wells in the mold. We then pipetted 60 µL of a 35% PAA solution (without VLPs, which forms the backing layer) on top of the first layer, followed by another centrifugation using the above parameters. This backing layer provides structural support to the MNs (**Figure 1D**), especially during their insertion into the skin. We used a positive displacement pipette (Rainin model MR-100) to add the viscous PAA solution to the PDMS mold.

**Figure 1:**
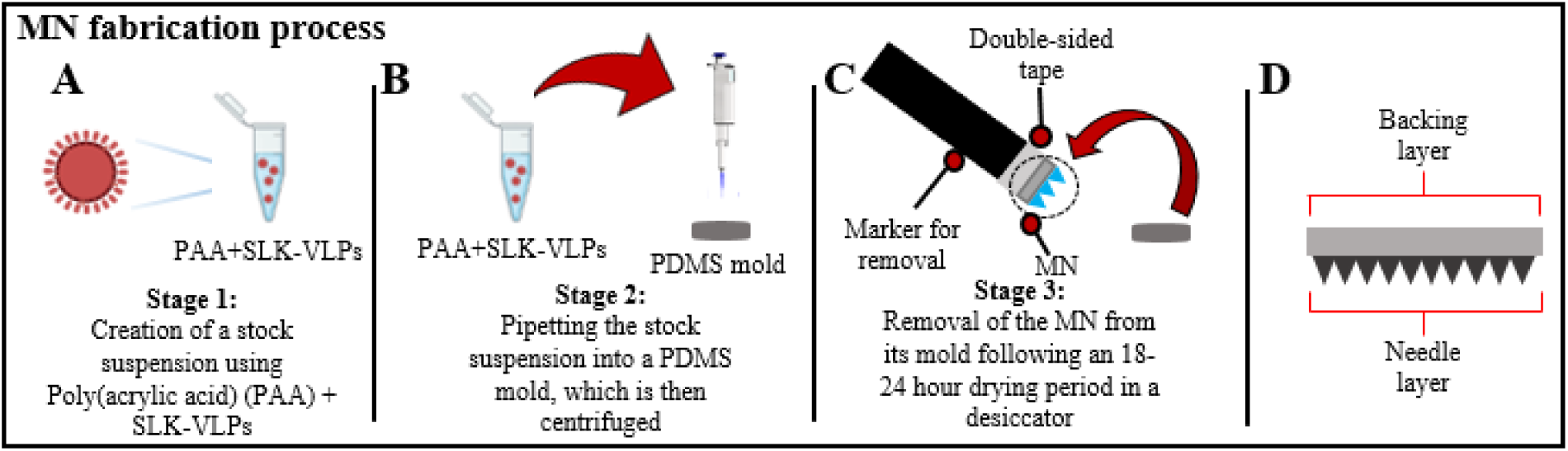
The 3 stages of MN fabrication (A-C) containing SLK-VLPs, and the resulting MN from the process (D). Images created using Biorender.

In the final stage, the MNs were dried in a desiccator for 18 to 24 hours. After drying, the MNs were removed from their molds using a flat-base device (in our case, a marker) attached to double-sided tape (**Figure 1C**). It was crucial to remove the MNs from the molds at a 90°C angle to prevent bending or breaking the needles.

### Needle Sharpness

An inverted fluorescent microscope (Olympus IX83, Tokyo, Japan) and the CellSens Dimensions software were used to assess needle sharpness and structural integrity of the MNs after fabrication. Images were taken using bright-field and fluorescence (FITC) imaging.

### Assessing VLP Conformation

We used the transmission electron microscope (TEM) to study the integrity of VLPs after loading them into MNs and compared them with pure SLK-VLPs (at similar VLP concentrations). We made VLP-loaded MNs without a backing layer for TEM imaging to minimize interference from PAA polymer. These MNs were dissolved in PBS before imaging the VLPs.

### Assessing the Particle Size Distribution of VLP-loaded MNs

The integrity of VLPs and their potential aggregation were investigated using Dynamic Light Scattering (DLS; Malvern Zetasizer Nano ZS, Malvern Instruments, Worcestershire, UK). We analyzed three different samples: pure VLPs (pre-fabrication stage), VLPs mixed with PAA solution (fabrication stage, before casting into PDMS molds), and VLP-containing MNs dissolved in PBS (post-fabrication stage). These samples showed varying degrees of VLP integrity and aggregation occurring at different stages of MN preparation. The DLS parameters included setting a temperature setting of 35°C and an equilibration time of 120 seconds.

### Microneedle Penetration Testing

To ensure our MNs could effectively penetrate the epithelial skin barrier, we created a skin mimic using lab-grade parafilm (Amcor; thickness-130µm). This involved folding the parafilm into an accordion shape with seven layers, resulting in a total thickness of 910µm (7 layers X130 µm each). A similar setup has been previously used to assess MN strength and penetration due to its affordability and similarity to porcine skin^2^. We inserted the MN into the parafilm using thumb pressure, analyzing the depth of needle penetration across the parafilm layers. A note of caution is that the parafilm lacks the elasticity and fluidity of mammalian skin, and this study only shows the MNs’ ability to breach a thick layer without losing needle integrity. Additionally, we tested MN penetration in freshly excised mouse skin (from the dorsal region) and confirmed needle penetration by observing fenestrations in the skin. While the thickness of mouse skin differs from that of humans, the penetration properties are comparable across the mammalian species^25^.

### Penetration Force

We studied the penetration force of MNs into excised mouse skin using a texture analyzer (TA; Stable Micro Systems TA.XTplus 100C). Blank MNs (without VLPs) were secured in place using an adaptor clamped inside a vertically moving probe (**Figure 2**). The excised mouse skin was gently stretched and secured over a holding chamber beneath the probe. Following this setup, the probe containing the MNs was lowered at a speed of 1 mm/sec towards the skin. We also evaluated blunted MNs (needles that have been filed down) and a 29G hypodermic needle as controls for comparison with intact MNs (**Figures 2 B and C**). Exponent Connect (version 8.0.11.0) software was used to visualize MN penetration into the skin, allowing us to calculate the penetration force.

**Figure 2:**
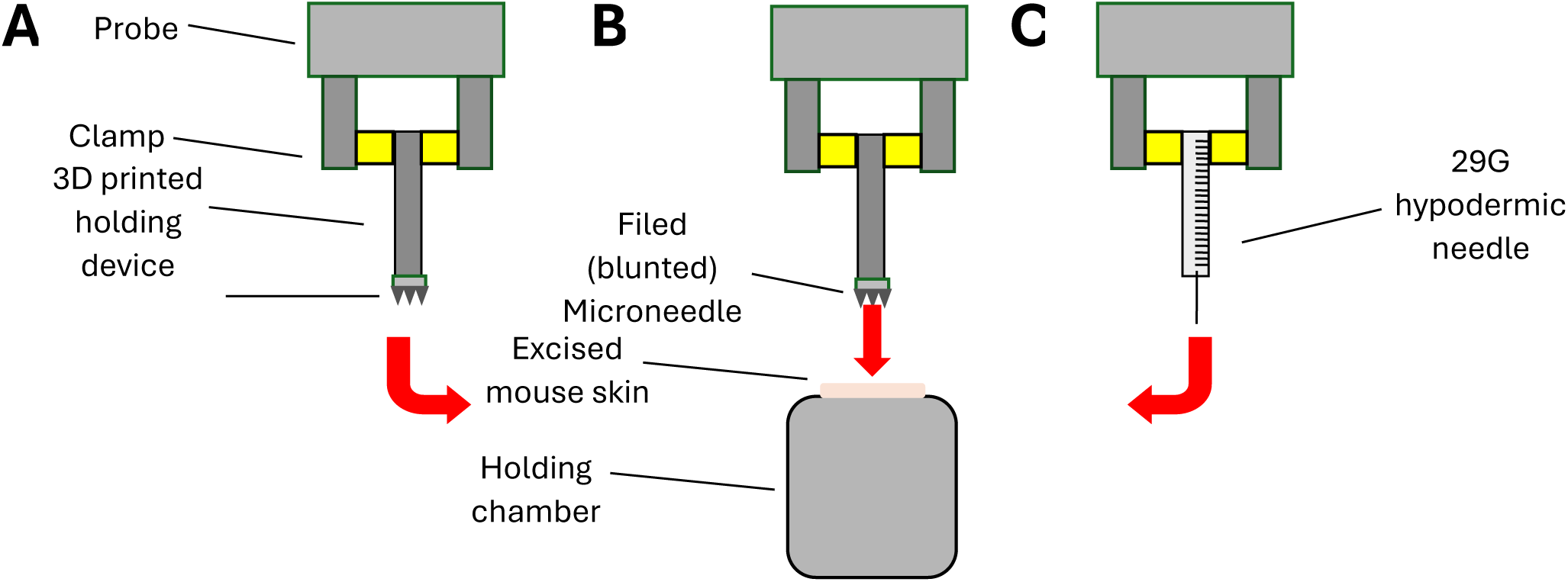
Study design for force penetration testing using (A) Intact MNs, (B) filed MNs, and (C) a hypodermic needle into excised mouse skin

### Dissolution Testing

To evaluate the dissolution of MNs in the interstitial fluid beneath the skin, we inserted MNs into freshly excised skin (dorsal region) from Balb/c mice. The MNs were removed at different times (up to 7 minutes post-insertion), and approximately 10µl of trypan blue dye (ThermoScientific, 0.2µm filtered) was added to highlight the micropores created in the skin. Excess dye was carefully removed from the skin before visualization, which was conducted using an iPhone 13.

This experiment was repeated to assess how the duration of applied pressure affects needle dissolution after insertion into the skin. MNs were inserted into the skin and held in place with thumb pressure for 30 seconds. They were then left in the skin without thumb pressure for different times (up to 5 minutes post-insertion). Light microscopy imaging was taken before and after MN insertion to evaluate needle dissolution.

### Thermostability Studies

MNs were stored at three different conditions-25°C, 40°C (in incubators), and 2°C (in a refrigerator) for approximately 140 days (5 months) to study the effects of varied and extended storage conditions on vaccine stability. Pure VLPs (not loaded in MNs) were also kept at these conditions for the same duration. The 40°C temperature represents extreme conditions observed in various regions across South America, Africa, and Asia, where many mosquito-borne diseases are endemic and the vaccine cold chain infrastructure is unreliable. The 25°C was chosen as it represents the global average temperature (room temperature), while 2-8°C is the standard temperature range for vaccine storage during transportation^26^. The MNs containing VLPs were formulated with and without backing layers to compare with freshly made VLP-MNs. We assessed parameters such as needle and VLP integrity, VLP aggregation, and VLP antigenicity. The MNs were kept in 12- and 24-well tissue culture plates, with individual MNs placed per well. The plates were sealed using either parafilm or duct tape.

### Immunizations in Mice (SLK-VLPs)

We evaluated the immunogenicity of MNs containing SLK-VLPs in 6- to 8-week-old female Balb/c mice (Jackson Laboratories). A total of 40 mice were immunized: 20 using SLK-MNs and 20 using 29G hypodermic needles for intramuscular (IM) injection (**Table I**). Each IM-injected mouse received a single dose of 10 µg VLPs. Twenty-four hours before immunization, the dorsal regions of the MN group were shaved. On the day of immunization, the mice were anesthetized with isoflurane. A single MN (10 µg VLP dose) was applied to the skin using thumb pressure to breach the skin and then secured in place with a clothespin for 10 minutes. After 10 minutes, the clothespin was removed, and the MN remained on the skin for an additional 10 minutes before being removed (totaling 20 minutes of application time). Sera were collected at 3, 6, 13, 18, and 24 weeks post-immunization.

**Table I.** Immunization conditions and total mice immunized using SLK-VLPs.

| Immunization Condition | SLK-MNs Freshly Made | SLK-MNs stored at 25°C | SLK-MNs stored at 40°C | SLK-VLPs Freshly Made (IM) | SLK-VLPs stored at 25°C (IM) | SLK-VLPs stored at 40°C (IM) | Total Number of Mice Immunized |
| --- | --- | --- | --- | --- | --- | --- | --- |
| Number of mice per group | 5 | 7 | 8 | 5 | 7 | 8 | 40 |

### Evaluating VLP Immunogenicity

Serum antibodies against SLK were detected using ELISA. Wells of Immulon2 ELISA plates (ThermoFisher Scientific) were coated with 500 ng of streptavidin overnight at 4°C. The following day, 250 ng of SMPH was added per well, and the plates were incubated for 1 hour at room temperature. Subsequently, wells were coated with 250 ng of SLK peptide and incubated for 2 hours at room temperature. Wells were blocked with PBS-0.5% milk for 1 hour at room temperature. Sera from immunized mice were serially diluted in PBS-0.5% milk and applied to wells overnight at 4°C. Reactivity to SLK was measured by adding HRP-labeled goat anti-mouse IgG (Jackson Immunoresearch), diluted 1:4000 in PBS-0.5% milk, and visualized by adding TMB substrate. The reaction was stopped using 1% HCl, and absorbance was measured at OD=450nm.

### Ethics statement for animal studies

All animal research was conducted in compliance with the Institutional Animal Care and Use Committee of the University of New Mexico School of Medicine (Approved protocol number: 22-201289-HSC).

### Statistical analysis

Statistical analysis was performed using Prism version 10 (GraphPad). A two-tailed t-test was used to compare the two groups. Significance was indicated when the value of p<0.05.

## Results

### Preparation of Microneedles

We formulated the MNs (**Figure 3**) containing VLPs using a three-stage process completed within 24 hours. The needles measured approximately 200 µm x 200 µm (base) and 600µm (height). Successful manufacturing was achieved by optimizing the addition of the PAA polymer (mixed with the VLPs) into the PDMS molds and the centrifugation parameters. The vaccine polymer mixture was pipetted using the MR-100 positive displacement pipette, which allowed for the complete ejection of the highly viscous solution into the molds, ensuring the accurate loading of the vaccine dose. The increased temperature setting (optimized to 30°C) during centrifugation facilitated a more efficient filling of the mold wells.

**Figure 3:**
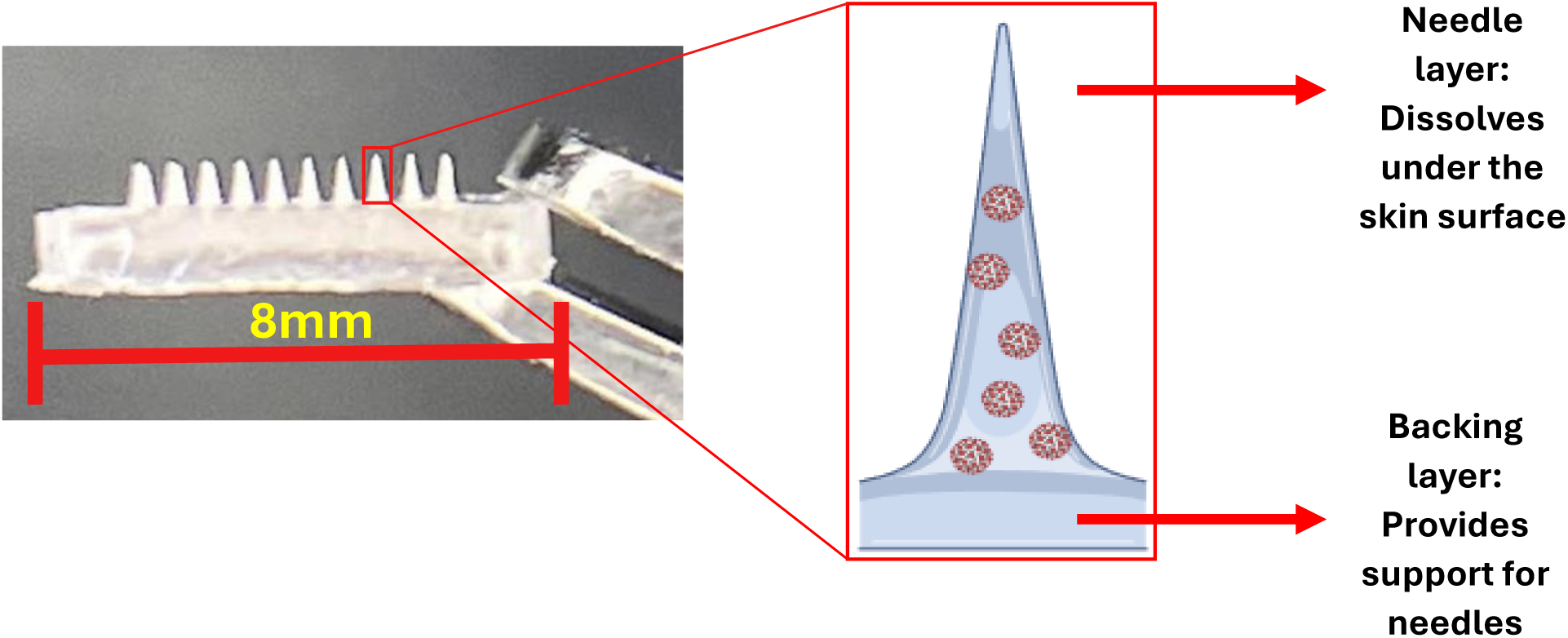
Post-fabrication MN containing VLPs. Created with Biorender.

### Characterization of Microneedles

#### MN Sharpness

The sharpness and structural integrity of the MNs are critical for effective vaccine delivery through the skin, as they need to breach the superficial layers, including the epidermis and dermis, which are rich in immune cells such as LCs and DCs. These cells play a vital role in detecting and capturing pathogens, thus initiating the host’s immune response mechanism. Light microscopy was employed to assess the sharpness and the number of intact needles. The MNs were imaged immediately after fabrication and after storage under different conditions (freshly made, stored at 25°C, and stored at 40°C for approximately 140 days). A small piece of orthodontic wax (adhesive) was placed on the edge of a glass slide to achieve the correct orientation of the MNs under the microscope (**Figure 4C**). The MN was placed onto the wax with the backing layer slightly angled downward, allowing for viewing successive rows of needles. As shown in **Figure 4 (A, B, D, and E)**, the needles maintained sharpness under all storage conditions, with over 90 intact needles remaining out of 100. Additionally, the needles exhibited pyramidal shape (**A and B**), which may enhance skin penetration compared to conical needles, as shown by others^27^; the penetration advantage of pyramidal needles likely stems from their greater surface area.

**Figure 4:**
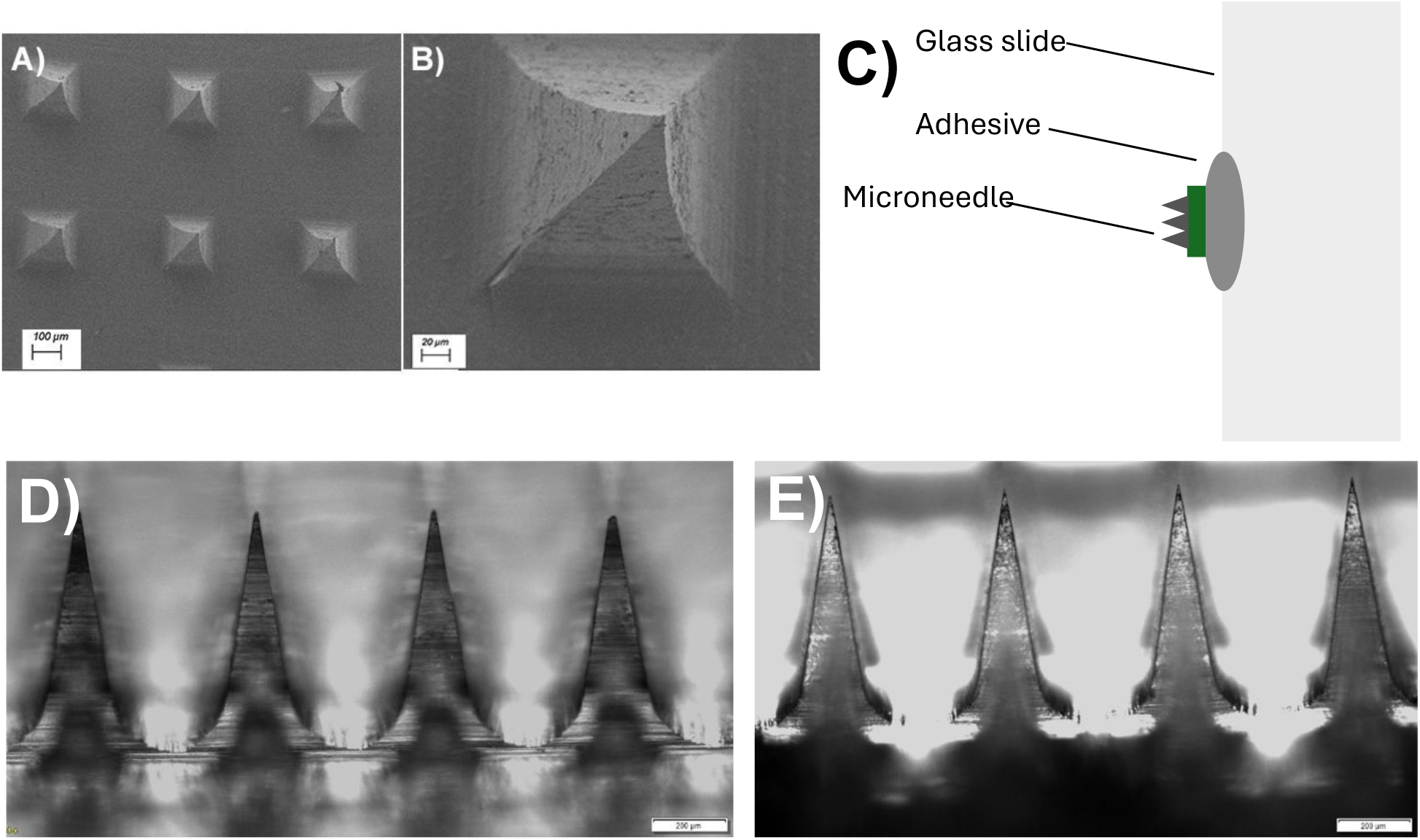
Assessment of MN sharpness and structural integrity after fabrication. (A,B) Scanning Electron Microscopy of MNs. (C) Schematic of MN placement onto a glass slide using an adhesive for imaging. (D) Image of freshly made MN. (E) Image of MN stored for ∼140 days at 40°C. Scale bars are shown in each panel.

The MNs kept at 2°C were deformed after removal from storage. Further testing revealed that MNs stored in high humidity (inside the refrigerator) led to deformed needles and decreased needle integrity. It is known that PAA polymer is hygroscopic due to the presence of polar functional groups that form hydrogen bonds with water molecules^28^. Therefore, we did not include MNs stored at 2°C for subsequent characterization and *in vivo* studies.

#### VLP aggregation

DLS was used to evaluate VLP aggregation at different stages of MN fabrication (pre-, during-, and post-fabrication steps). Control samples (pure VLPs in PBS) exhibited an unimodal particle size distribution (PSD) with a uniform size of approximately 30nm (**Figure 5A**). **Figure 5B** shows the PSD of VLPs mixed with PAA polymer (29%v/v), revealing a slight shift in the average size (∼25nm). In the post-fabrication sample (**Figure 5C**), the VLPs remained uniform and similar in size to the pure VLPs, indicating that their integrity was preserved after MN fabrication. Further, the absence of multimodal peaks suggests that VLP aggregation did not occur after loading and dissolving from the MNs.

**Figure 5:**
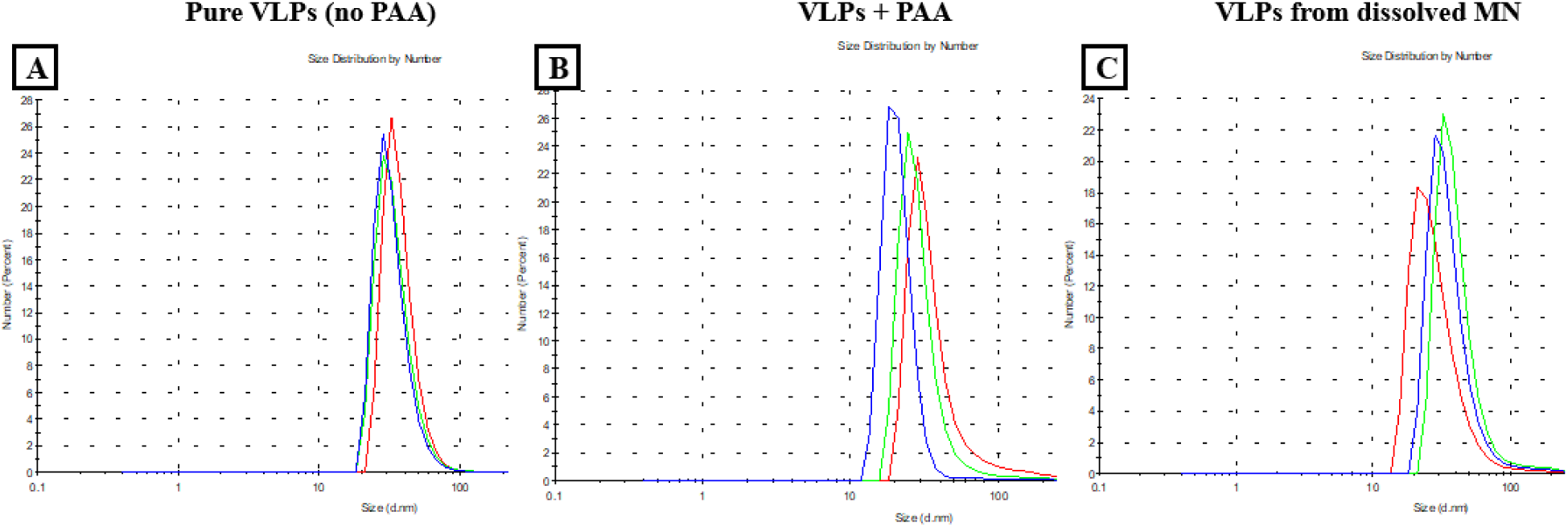
Particle Size Distribution (PSD) of the VLPs at different stages of fabrication. (A) Pure VLPs in PBS (no PAA). (B) VLPs mixed with 29% v/v PAA. (C) VLPs from a dissolved MN (post-fabrication).

#### Microneedle Penetration

We tested the penetration capacity of blank MNs using a seven-layered folded parafilm. The needles successfully penetrated four layers, each approximately 130µm thick. This study demonstrates that the needles penetrated approximately 520µm (**Figure 6**), corresponding to nearly the entire needle length, and confirms that the needles can breach a thick barrier without breaking (data not shown). It is important to note that this study does not serve as a substitute for skin penetration studies but only demonstrates proof-of-concept regarding the MN’s ability to penetrate a thick layer while maintaining needle integrity.

**Figure 6:**
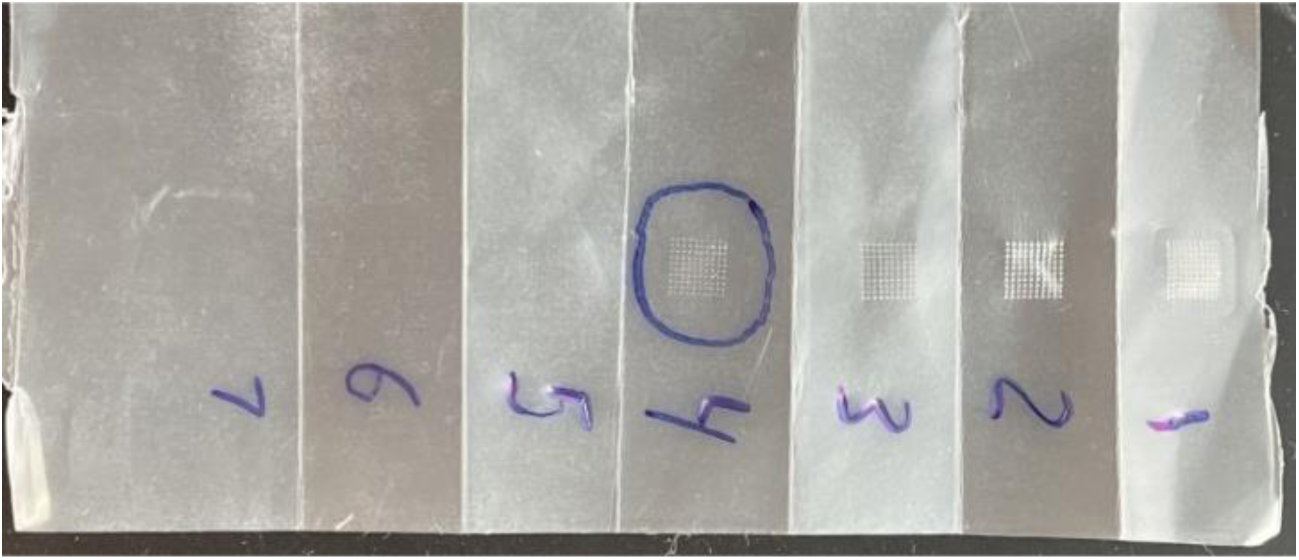
Parafilm folded accordion-style (7 layers) for assessment of MN penetration. The MN penetrated the fourth layer (>90% needles).

#### *Ex-vivo* Microneedle Penetration

We showed that VLP-containing MNs could penetrate excised mouse skin *ex vivo*. Trypan blue dye stained the pores created by the needles (**Figure 7**), showing skin pores (or fenestrations; **Panel B**) for all needle arrays (10X10).

**Figure 7:**
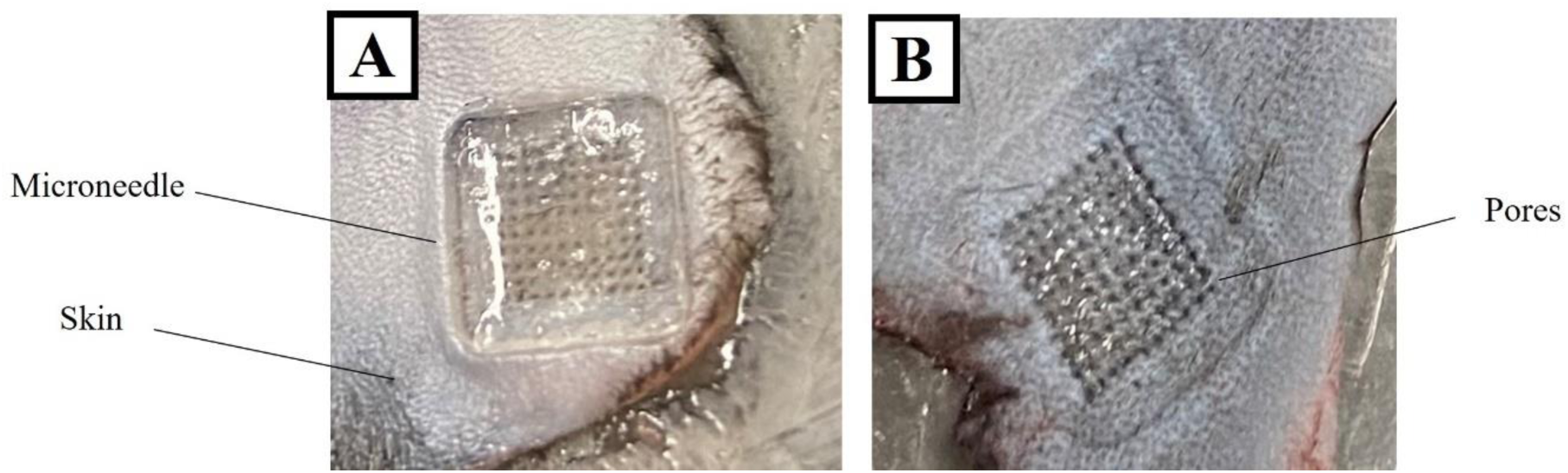
MN placed in excised mouse skin (A) After removal of the MN, trypan blue staining reveals fenestrations caused by the needle insertions.

#### Penetration Force

Breaching the skin barrier with vaccine-containing MNs is essential for successful vaccination. We evaluated the force at which the MNs (**Figure 8A, D**) could break through this skin barrier. We compared these results to a blunted MN (filed down; **Figure 8B, E**) and a 29G hypodermic needle (**Figure 8C**). The resulting data indicated that the intact MN exerted a skin-breaching force comparable to that of a traditional hypodermic needle (78mN and 73mN, respectively; **Figure 8F**). However, the blunted (filed) MN exhibited a higher penetrating force of 210mN, as expected, since these needles are not sharp and, therefore, have difficulty breaching the skin barrier compared to the intact MN and the hypodermic needle.

**Figure 8:**
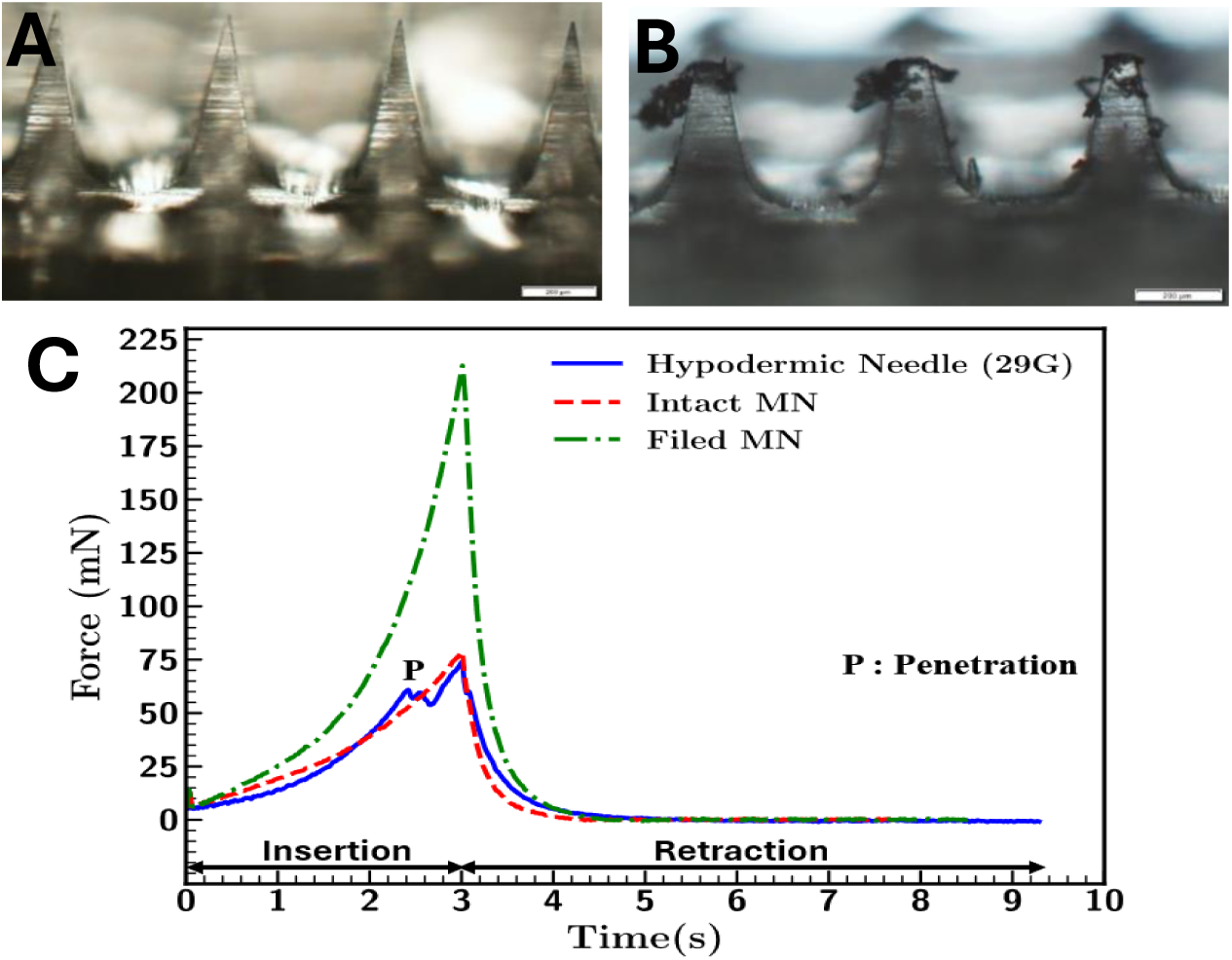
Penetration force required for vaccination. Representative images of (A) Intact and (B) blunted MNs used for force insertion assessment in excised mouse skin (scale bars-200µm). (C) Comparison of penetration force required for intact MN (red), blunted MN (green), and hypodermic needle (blue) (C).

The penetration force exerted by the intact MN was 78mN, which agrees with previously reported experimental tests on mouse skin^29^ and finite element modeling^30^. This result is consistent with findings from Zhang et al., who reported that dissolvable MNs exhibit a penetration force of 89mN in mouse skin^31^. Conversely, dissolvable MNs require a significantly greater insertion force of approximately 30N in pig skin^29^. This difference is likely due to Young’s modulus (*E*) of the two skin types (mouse versus pig), which measures material stiffness and the material’s ability to deform. Pig skin (*E* = 7.34 kPa) is stiffer than mouse skin (*E* = 3.81 kPa)^25^, indicating that our MNs would likely require more force to penetrate human skin, which tends to have even greater stiffness depending on variable factors such as pediatric versus geriatric skin.

#### Structural Integrity

TEM was used to evaluate the integrity of VLP following their incorporation into MNs. Before imaging, MNs (without backing layer) containing VLPs were dissolved in PBS and compared to pure VLPs at similar concentrations. As shown in **Figures 9A and B**, the integrity and concentration of the VLP remained intact throughout the MN fabrication process. The VLP size was approximately 30nm, consistent with our previous findings^5^ and those shown in **Figure 5 (DLS)**.

**Figure 9:**
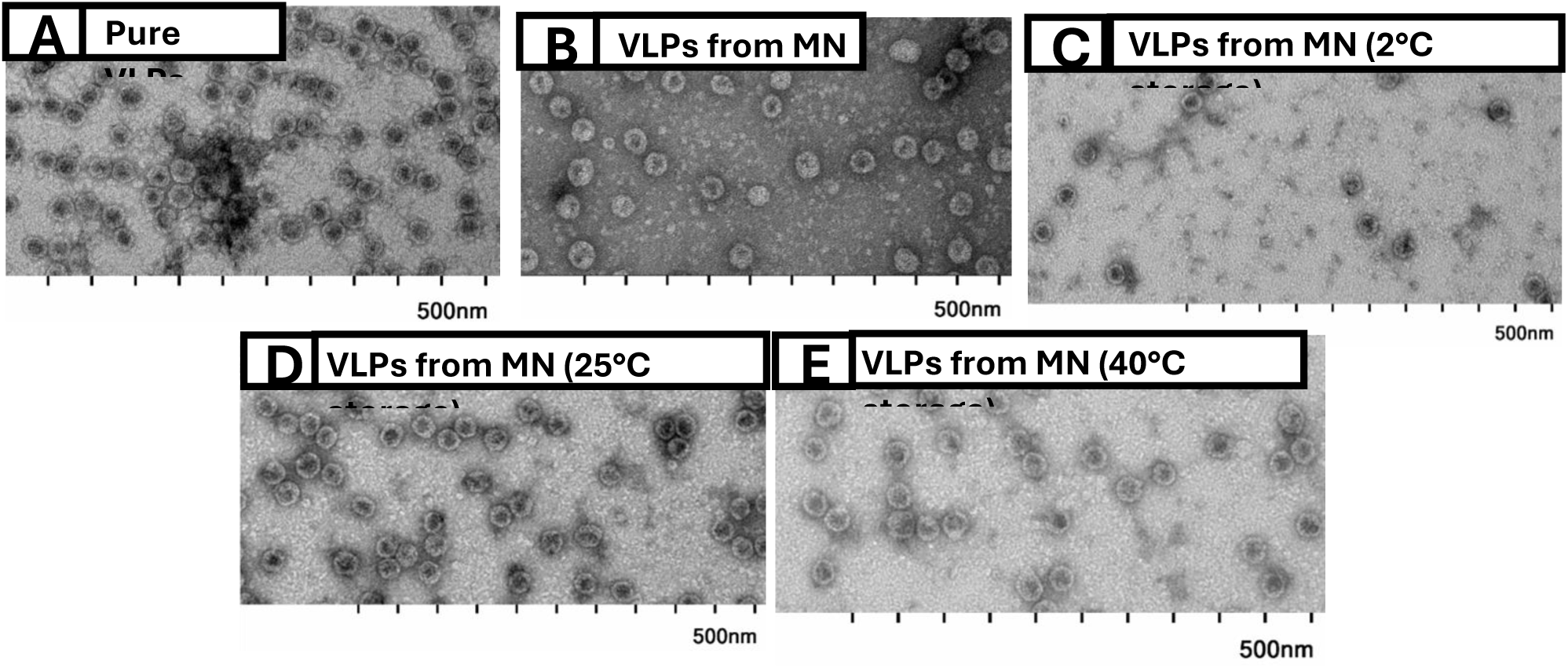
TEM imaging of SLK-VLPs. (A) Pure VLPs in PBS (no PAA) and (B) VLPs from freshly made MNs. VLPs were stored in MNs at (C) 2°C, (D) 25°C, and (E) 40°C for ∼140 days.

We also evaluated the integrity of the VLPs after storing the MNs for approximately 140 days under three different temperature conditions (**Figure 9C-E**): 2°C, 25°C, and 40°C. VLPs in MNs stored at both 25°C and 40°C maintained their integrity during extended storage, indicating the stability of the VLPs within the MNs. Additionally, despite some needle deformation observed in the MNs stored at 2°C, the VLPs retained their integrity. The robustness of our MNs and the manufacturing process in preserving VLP integrity against temperature exposures can improve their shelf life, simplifying the vaccine’s storage, transport, and distribution without compromising its stability.

#### Dissolution Testing

We studied the dissolution rate of MNs after they were inserted into freshly excised mouse skin for 1, 3, and 5 minutes. The dissolution of MNs in the interstitial fluid and the subsequent release of VLPs are critical for eliciting a robust immune response, as this region contains immune cells that play a crucial role in immune surveillance against pathogens. We investigated MN dissolution using two different methods. In the first study, individual MNs were inserted in the skin using thumb pressure only and were kept embedded for 1, 3, and 5 minutes. After MN removal, the skin was imaged using light microscopy. We observed a reduction in needle length by approximately 50%, 80%, and 99%, respectively (**Figure 10**).

**Figure 10:**
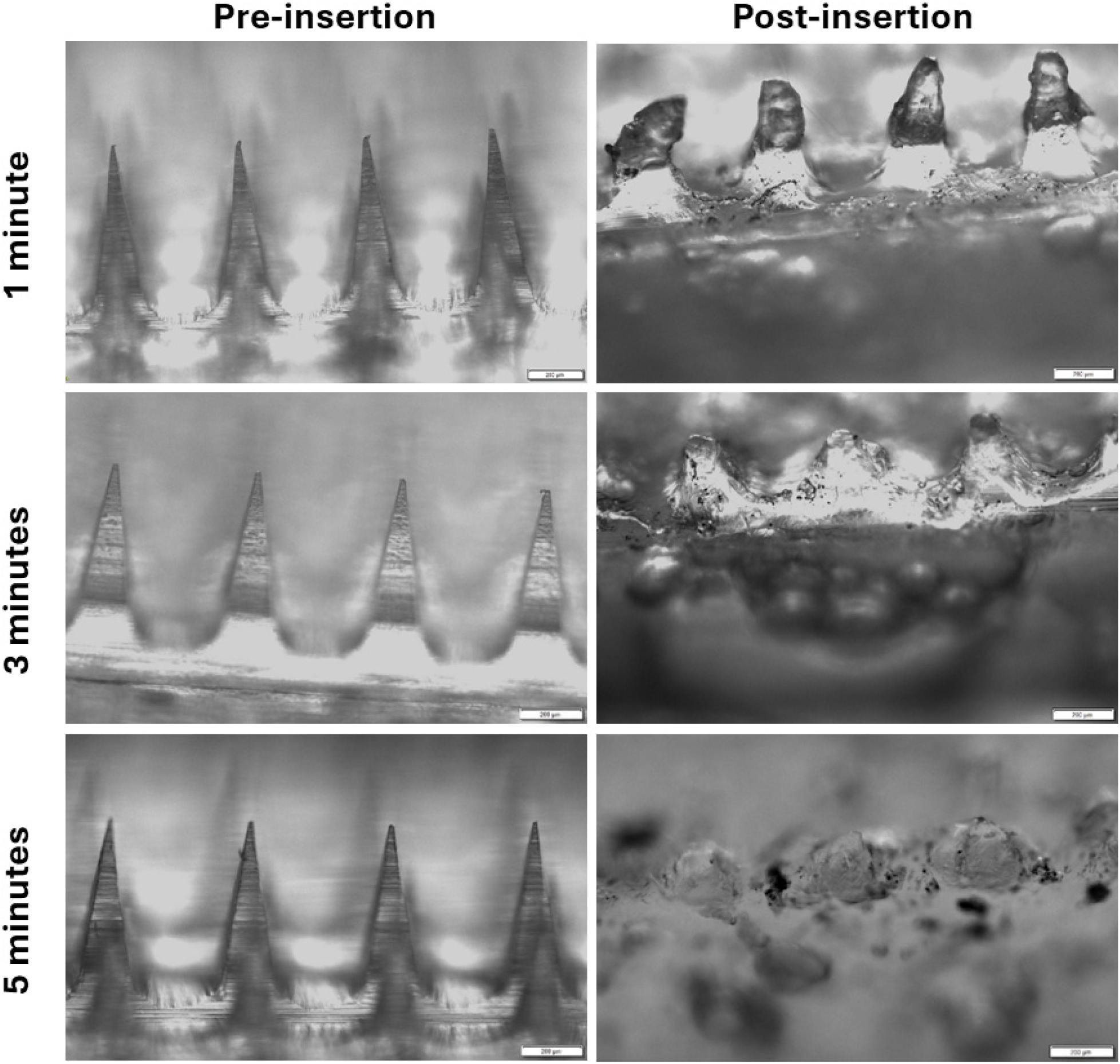
Images of needles before and after inserting MNs into excised mouse skin with constant thumb pressure for 1, 3, and 5 minutes.

The second dissolution study involved inserting the MN with thumb pressure, maintaining the pressure on the skin for 30 seconds, and then releasing the pressure. The MNs were then left on the skin for different time intervals: 2.5 minutes and 4.5 minutes, totaling 3 and 5 minutes, respectively. Similar to the first study, more needles dissolved under the skin with increased time (**Figure 11**).

**Figure 11:**
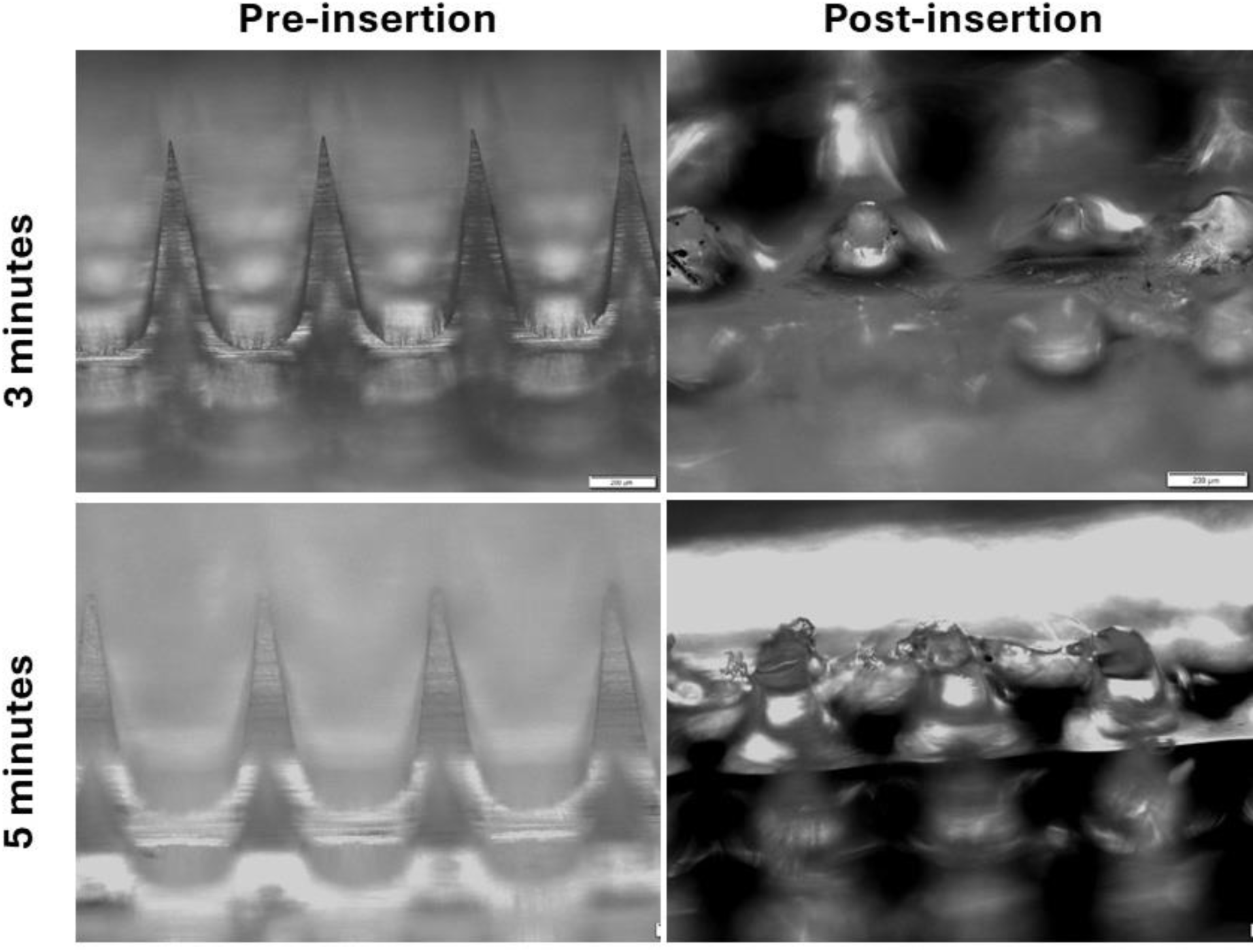
Images of needles before and after inserting MNs into excised mouse skin without constant thumb pressure for 3 and 5 minutes.

#### Heat-stable microneedles induce high antibody titers

We assessed IgG antibody titers in blood from mice after they were vaccinated with a single dose of SLK-VLPs via hypodermic needles (IM; intramuscular) and MNs. The mice received vaccines that were either freshly prepared or stored at 25°C or 40°C for approximately 140 days (Table 1). Sera were collected at regular intervals (3, 6, 8, 12, 18, and 24 weeks post-immunization) to evaluate anti-SLK IgG antibody titers. Both MN and IM vaccination led to strong and durable anti-SLK antibody responses for 6 months following vaccination, at which point the mice were sacrificed (**Figure 12**). The antibody titers remained consistent across all vaccinated groups during this period; therefore, only the titers at 6 months are presented. There was no significant difference when comparing either the IM or MN groups under the three tested conditions. However, the titers for both freshly prepared IM and 25°C IM groups were significantly higher compared to those for the 40°C IM and MN groups (p<0.05). No significant difference was observed among the MN groups.

**Figure 12:**
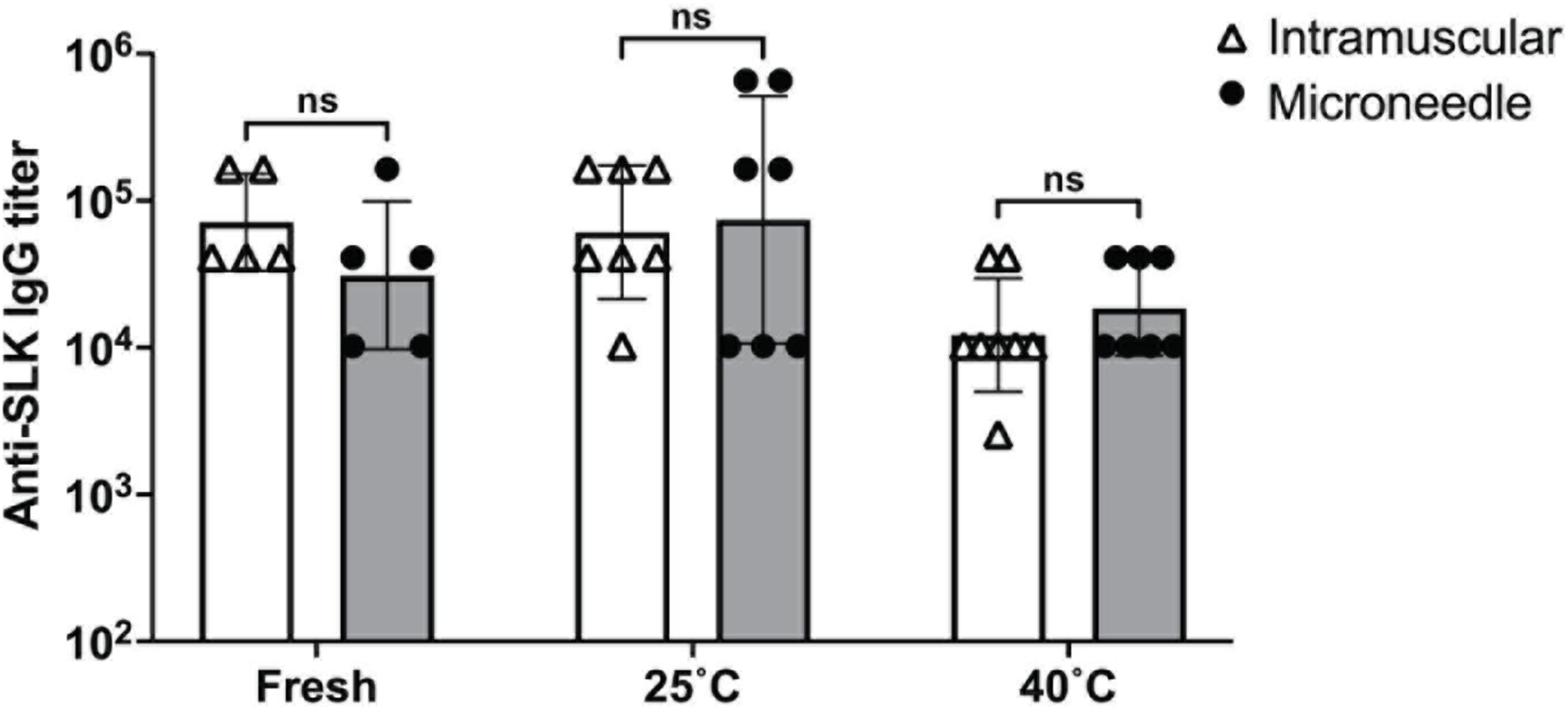
MNs containing VLPs elicit high-titer and long-lasting antibody responses against SLK. Balb/c mice were immunized with a single (10µg) dose of SLK VLPs delivered via hypodermic needle (IM) or MNs that were (A) made fresh (n=5), (B) stored at 25°C for ∼140 days (n=7), or (C) stored at 40°C for ∼140 days (n=8). Geometric mean IgG titers were determined longitudinally over 24 weeks after immunization. Data is represented as geometric mean ± geometric SD. Statistical significance was determined by a two-tailed t-test (p<0.05).

Mice immunized with MNs exhibited slightly elevated titers in the 25°C and 40°C groups compared to their IM counterparts, suggesting that the SLK-VLP vaccine (in liquid) retains its integrity at elevated temperatures. Additionally, loading the vaccine into MNs did not affect its integrity. The 40°C group displayed the lowest antibody titers overall at the 6-month timepoint, while the 25°C group had the highest overall titer count at the same time. However, the titers across all groups were relatively similar (ranging from 10^4^-10^5^) and were not significantly different.

## Discussion

Novel immunization strategies will play a crucial role in future immunizations by addressing the challenges faced during the COVID-19 pandemic, including the reliance on needles and syringes for vaccine administration and the weakness in vaccine distribution supply chains. Vaccine production costs are high, even in high- and middle-income countries (HMICs)^32^, where most vaccine development and manufacturing occur. Due to the increasing global population and a lack of robust supply chain networks, LMICs will continue to face challenges accessing vaccines, especially in remote regions. However, LMICs could benefit from developing their own vaccine manufacturing capabilities, lowering production costs, and contributing to economic development. Emerging epidemics will drive the need for novel vaccine delivery strategies in both HMICs and LMICs.

Our research shows that MNs containing VLPs remain stable for up to five months under high-temperature storage conditions. Thermostable vaccines would also enhance global pandemic preparedness by ensuring that vaccines can be developed, stockpiled, and deployed quickly in the event of another global health crisis. Typically, the fabricating process for dissolvable MNs processes using micromolding requires longer drying times^33^; for example, dissolving MNs containing an influenza vaccine required at least two days for fabrication^34^. Our MNs can be fabricated within 24 hours using a safe and biocompatible polymer. A simplified and faster vaccine manufacturing process will enable rapid vaccination efforts in response to new or emerging infectious disease outbreaks.

Characterization of our VLP-MNs revealed that they are strong and capable of penetrating the mouse skin to deliver the vaccine into the dermal region. We also demonstrated that the needles do not break upon insertion into a skin mimic (folded parafilm) when inserted with thumb pressure. The MNs dissolve beneath the skin surface (*ex vivo*) within five minutes after insertion, allowing for the immediate release of the vaccine contents. Additionally, the insertion force required for MNs into mouse skin (*ex vivo*) is comparable to a traditional hypodermic needle.

Our study showed the integrity of the vaccine (VLP) is maintained during MN manufacture and after storing the MNs at elevated temperatures for extended periods. We did not observe any clumping or aggregation of the VLPs after MN reconstitution, either in freshly made MNs or stored MNs.

Our MNs were loaded with VLPs conjugated to SLK, a salivary peptide that facilitates arbovirus transmission by *Aedes aegypti* mosquitoes. Arboviral infections have emerged as significant public health concerns, with common infections including dengue (*Flavivirus*), chikungunya (*Alphavirus*)^35^ and Zika (*Flavivirus*)^36^. A 2013 study showed that dengue’s global burden is nearly 400 million infections every year^37^. Despite the diversity of arboviral diseases, the transmission mechanism from mosquitoes to humans remains the same. After biting through the skin of a mammalian host, the infected mosquito releases its saliva containing the virus. SLK, a vasoactive salivary protein, increases host susceptibility by enhancing blood vessel permeability and edema, allowing for a greater influx of virus-permissive cells and higher viral load, which can lead to more severe disease outcomes^38^. We anticipate that immunizing beneath the skin surface with MNs, where arboviruses are first transmitted, would elicit a local immune response due to the abundance of APCs in that region^39^.

Our findings highlight the robustness of our dissolvable MNs in maintaining the integrity and stability of VLPs. We demonstrate that MN administration elicits robust immune responses in mice comparable to IM immunization. Further, *in vivo* immunogenicity of the VLP-MNs is not affected by storage at extreme temperatures for extended periods. Our previous studies indicated that VLPs formulated with stabilizing excipients maintain their potency under elevated temperature conditions over long storage periods^5^.

We show that dissolvable VLP-MNs are comparable to traditional needle vaccination methods in terms of immune response generation. However, MNs offer significant benefits over IM or subcutaneous (SC) injections using hypodermic needles (**Table II**). After administration, MNs do not engage the pain receptors, making them minimally invasive. Additionally, MNs allow for more straightforward delivery since the vaccine is pre-loaded. In contrast, hypodermic needles require the extra step of drawing the (liquid) vaccine from a vial into a syringe before administration. Moreover, the region beneath the skin, where the vaccine is delivered using MNs, has more immune cells than the muscle (IM route) or adipose tissue (SC route)^40^.

**Table II.** Comparing needle administration techniques.

| Comparing Needle Administration Techniques for Vaccination |  |  |
| --- | --- | --- |
|  | Traditional vaccine administration by needle and syringe | Dissolvable microneedle administration |
| **Route of administration** | Intramuscular or subcutaneous<br>• Extends into or below subcutaneous tissue<br>• More likely to interact with pain receptors | Transdermal<br>• Extends through the epidermis and possibly part of the dermis<br>• Little to no interaction with pain receptors |
| **Method of administration** | Carefully extract and inject the correct vaccine dose into the appropriate site.<br>• Requires trained staff<br>• It can lead to needlestick injuries if done improperly<br>• Large space required for vaccine vials, sterile needles, and syringes | Place MNs on the skin, apply pressure to insert, and allow time for needles to dissolve.<br>• Simple, and can potentially be administered without training.<br>• Space saving allows more doses to be stored. |
| **Uses** | Designed strictly for injecting vaccine | Capable of both storing and administering vaccine |
| **Injection site characteristics** | Intramuscular<br>• Low immune cell density<sup>40,41</sup><br>• High blood supply<br>Subcutaneous<br>• Low immune cell density<sup>40</sup><br>• High blood supply | Transdermal<br>• Higher immune cell density<sup>40</sup><br>• Limited blood supply |

However, MNs as a vaccine delivery platform have certain limitations. First, the amount of vaccine that can be loaded in the MNs is limited; a high vaccine dose in small MNs used here (8 × 8 mm) would require a reduction in polymer content, which could compromise the strength of the needles, increasing the risk of breakage during insertion. Our study showed that mice developed a robust immune response after MNs administration, comparable to IM vaccination. However, we used a relatively small vaccine dose (10µg) in the MNs and for IM injection that achieved adequate *in vivo* immunogenicity. This dose limitation, particularly with smaller MNs, could be addressed using multiple smaller MNs, single larger MNs, or providing vaccine booster doses. Another challenge we encountered was the deformation of MNs when stored in the refrigerator under high humidity conditions. This issue arose due to the hygroscopic nature of the PAA polymer used to make our MNs^28^. We can mitigate this concern by storing PAA-based MNs in moisture-protective packaging^42^ or opting for alternative polymers for MN fabrication.

In our previous research, two VLP doses were necessary to generate robust and long-term immunity in mice^43^. However, in this study, mice were immunized with a single VLP dose (both IM and MNs), which elicited a strong and sustained immune response systemically. Future studies should focus on determining the optimal vaccine dose and number of booster doses needed to maintain long-term immunity with MN-delivered vaccines, which was not explored in this study.

To effectively immunize against arboviruses (or any other disease using MN administration) on a large scale, MN manufacturing processes must be compatible with large-scale production. Currently, the 24-hour MN manufacturing process described here serves as a proof-of-concept study demonstrating lab-scale capability for smaller batch sizes. Our MN fabrication involved manual steps, including pipetting polymer-VLP suspension into PDMS molds, centrifuging, and transferring the molds to drying chambers. Automating this process would enable industry-scale manufacturing, ultimately facilitating MN commercialization and mass immunization. Several companies worldwide have developed automated capabilities for MN manufacturing.

In addition to ensuring good manufacturing practices for reproducible and scalable MN production, it is essential to confirm the safety and biocompatibility of the excipients (including polymers) to prevent infections or unwanted immune responses that could interfere with the vaccine’s efficacy. These safety studies must be conducted in human trials before MNs can be approved by the regulatory agencies for commercial use. A phase 1 clinical trial of dissolving MNs targeting influenza reported strong antibody responses comparable to those achieved through IM immunization^44^. Recently, the use of MNs for delivering biologics in therapeutic applications has also garnered interest^45^. Furthermore, MNs have improved patient acceptance among diabetic patients by minimizing the pain associated with SC insulin injection in both pediatric and adult patients. A phase 2/3 clinical trial in type 1 diabetic patients indicated a faster onset and offset of insulin effects using (hollow) MNs compared to traditional SC injections^46^.

## Conclusion

We have demonstrated a lab-scale manufacturing process for PAA-based MNs loaded with VLPs conjugated to an *Aedes aegypti* salivary peptide (SLK). Thermostable vaccines are essential for mass vaccination efforts, especially in LMICs, where maintaining a robust cold-chain infrastructure can be challenging. The MNs displayed a similar skin insertion force to that of traditional hypodermic needles used for IM injections, and they successfully penetrated the skin and dissolved quickly to release the vaccine. Finally, the VLP-MNs exhibited comparable immunogenicity to the IM-administered vaccine even after being stored at elevated temperatures for approximately five months.

## Acknowledgments

We acknowledge Tamara Howard and the UNM Electron Microscopy Facility for supporting TEM imaging. We also acknowledge Yogesh Nepal, Lauren Crowley, Benjamin Marwedel, and Andrea Ibarra for assisting with the animal studies and collecting pertinent DLS data. We also acknowledge facilities provided by the Autophagy, Inflammation, & Metabolism (AIM) Center of Biomedical Research Excellence (COBRE) cores, funded by NIH grant P20 GM121176.

## Author Contributions

A.L. helped design and execute all aspects of the project, including the preparation, storage, and characterization of the microneedle, and contributed equally to primary authorship. A.F. prepared VLPs for microneedle application, performed and coordinated animal studies, generated and analyzed immune data, and also contributed equally to primary authorship. M.R. assisted in designing microneedle preparation and characterization studies and helped edit the paper. A.B. helped to create a protocol for microneedle preparation. R.J. coordinated the texture analyzer studies and contributed to writing specific sections of the paper. N.J. assisted in editing the manuscript and provided expertise on strength and insertion studies related to the texture analyzer. B.C. contributed to the writing of the paper and offered expertise on VLP manufacture and preclinical immune studies following vaccination. P.M. helped plan and coordinate all project studies, analyze results, and contribute to the paper’s writing and editing.

## REFERENCES

(1) Carter, A.; Msemburi, W.; Sim, S. Y.; Gaythorpe, K. A. M.; Lambach, P.; Lindstrand, A.; Hutubessy, R. Modeling the Impact of Vaccination for the Immunization Agenda 2030: Deaths Averted Due to Vaccination against 14 Pathogens in 194 Countries from 2021 to 2030. Vaccine 2023. 10.1016/j.vaccine.2023.07.033.

(2) Smallpox | CDC. https://www.cdc.gov/smallpox/index.html (accessed 2023-07-28).

(3) Kunda, N. K.; Wafula, D.; Tram, M.; Wu, T. H.; Muttil, P. A Stable Live Bacterial Vaccine. Eur. J. Pharm. Biopharm. 2016, 103, 109–117. 10.1016/j.ejpb.2016.03.027.

(4) Price, D. N.; Kunda, N. K.; Ellis, R.; Muttil, P. Design and Optimization of a Temperature-Stable Dry Powder BCG Vaccine. Pharm. Res. 2019, 37 (1), 11. 10.1007/s11095-019-2739-8.

(5) Saboo, S.; Tumban, E.; Peabody, J.; Wafula, D.; Peabody, D. S.; Chackerian, B.; Muttil, P. Optimized Formulation of a Thermostable Spray-Dried Virus-Like Particle Vaccine against Human Papillomavirus. Mol. Pharm. 2016, 13 (5), 1646–1655. 10.1021/acs.molpharmaceut.6b00072.

(6) Yarlagadda, H.; Patel, M. A.; Gupta, V.; Bansal, T.; Upadhyay, S.; Shaheen, N.; Jain, R. COVID-19 Vaccine Challenges in Developing and Developed Countries. Cureus 14 (4), e23951. 10.7759/cureus.23951.

(7) Vaz, K.; McGrowder, D.; Alexander-Lindo, R.; Gordon, L.; Brown, P.; Irving, R. Knowledge, Awareness and Compliance with Universal Precautions among Health Care Workers at the University Hospital of the West Indies, Jamaica. 2010, 1 (4).

(8) Motaarefi, H.; Mahmoudi, H.; Mohammadi, E.; Hasanpour-Dehkordi, A. Factors Associated with Needlestick Injuries in Health Care Occupations: A Systematic Review. J. Clin. Diagn. Res. JCDR 2016, 10 (8), IE01–IE04. 10.7860/JCDR/2016/17973.8221.

(9) Love, A. S.; Love, R. J. Considering Needle Phobia among Adult Patients During Mass COVID-19 Vaccinations. J. Prim. Care Community Health 2021, 12, 21501327211007393. 10.1177/21501327211007393.

(10) Ashok, A.; Brison, M.; LeTallec, Y. Improving Cold Chain Systems: Challenges and Solutions. Vaccine 2017, 35 (17), 2217–2223. 10.1016/j.vaccine.2016.08.045.

(11) Weniger, B. G.; Papania, M. J. Alternative Vaccine Delivery Methods. Vaccines 2013, 1200–1231. 10.1016/B978-1-4557-0090-5.00063-X.

(12) Kim, Y.-C.; Park, J.-H.; Prausnitz, M. R. Microneedles for Drug and Vaccine Delivery. Adv. Drug Deliv. Rev. 2012, 64 (14), 1547–1568. 10.1016/j.addr.2012.04.005.

(13) Preston, K. B.; Randolph, T. W. Stability of Lyophilized and Spray Dried Vaccine Formulations. Adv. Drug Deliv. Rev. 2021, 171, 50–61. 10.1016/j.addr.2021.01.016.

(14) Mistilis, M. J.; Bommarius, A. S.; Prausnitz, M. R. Development of a Thermostable Microneedle Patch for Influenza Vaccination. J. Pharm. Sci. 2015, 104 (2), 740–749. 10.1002/jps.24283.

(15) Connelly, D. Microneedles: a new way to deliver vaccines. The Pharmaceutical Journal. https://pharmaceutical-journal.com/article/feature/microneedles-a-new-way-to-deliver-vaccines (accessed 2023-06-21).

(16) Donnelly, R. F.; Singh, T. R. R.; Garland, M. J.; Migalska, K.; Majithiya, R.; McCrudden, C. M.; Kole, P. L.; Mahmood, T. M. T.; McCarthy, H. O.; Woolfson, A. D. Hydrogel-Forming Microneedle Arrays for Enhanced Transdermal Drug Delivery. Adv. Funct. Mater. 2012, 22 (23), 4879–4890. 10.1002/adfm.201200864.

(17) Moore, L. E.; Vucen, S.; Moore, A. C. Trends in Drug- and Vaccine-Based Dissolvable Microneedle Materials and Methods of Fabrication. Eur. J. Pharm. Biopharm. 2022, 173, 54–72. 10.1016/j.ejpb.2022.02.013.

(18) Ma, G.; Wu, C. Microneedle, Bio-Microneedle and Bio-Inspired Microneedle: A Review. J. Controlled Release 2017, 251, 11–23. 10.1016/j.jconrel.2017.02.011.

(19) Health, C. for D. and R. Regulatory Considerations for Microneedling Products. https://www.fda.gov/regulatory-information/search-fda-guidance-documents/regulatory-considerations-microneedling-products (accessed 2024-09-30).

(20) West, H. C.; Bennett, C. L. Redefining the Role of Langerhans Cells As Immune Regulators within the Skin. Front. Immunol. 2018, 8.

(21) Zaric, M.; Lyubomska, O.; Poux, C.; Hanna, M. L.; McCrudden, M. T.; Malissen, B.; Ingram, R. J.; Power, U. F.; Scott, C. J.; Donnelly, R. F.; Kissenpfennig, A. Dissolving Microneedle Delivery of Nanoparticle-Encapsulated Antigen Elicits Efficient Cross-Priming and Th1 Immune Responses by Murine Langerhans Cells. J. Invest. Dermatol. 2015, 135 (2), 425–434. 10.1038/jid.2014.415.

(22) Tumban, E.; Peabody, J.; Peabody, D. S.; Chackerian, B. A Pan-HPV Vaccine Based on Bacteriophage PP7 VLPs Displaying Broadly Cross-Neutralizing Epitopes from the HPV Minor Capsid Protein, L2. PloS One 2011, 6 (8), e23310. 10.1371/journal.pone.0023310.

(23) Aida, Y.; Pabst, M. J. Removal of Endotoxin from Protein Solutions by Phase Separation Using Triton X-114. J. Immunol. Methods 1990, 132 (2), 191–195. 10.1016/0022-1759(90)90029-u.

(24) Larrañeta, E.; Moore, J.; Vicente-Pérez, E. M.; González-Vázquez, P.; Lutton, R.; Woolfson, A. D.; Donnelly, R. F. A Proposed Model Membrane and Test Method for Microneedle Insertion Studies. Int. J. Pharm. 2014, 472 (1), 65–73. 10.1016/j.ijpharm.2014.05.042.

(25) Wang, Y.; Marshall, K. L.; Baba, Y.; Gerling, G. J.; Lumpkin, E. A. Hyperelastic Material Properties of Mouse Skin under Compression. PLoS ONE 2013, 8 (6), e67439. 10.1371/journal.pone.0067439.

(26) ACIP Storage and Handling Guidelines for Immunization | CDC. https://www.cdc.gov/vaccines/hcp/acip-recs/general-recs/storage.html (accessed 2024-04-22).

(27) Lee, J. W.; Park, J.-H.; Prausnitz, M. R. Dissolving Microneedles for Transdermal Drug Delivery. Biomaterials 2008, 29 (13), 2113–2124.

(28) Arkaban, H.; Barani, M.; Akbarizadeh, M. R.; Pal Singh Chauhan, N.; Jadoun, S.; Dehghani Soltani, M.; Zarrintaj, P. Polyacrylic Acid Nanoplatforms: Antimicrobial, Tissue Engineering, and Cancer Theranostic Applications. Polymers 2022, 14 (6), 1259. 10.3390/polym14061259.

(29) Bhatnagar, S.; Bankar, N. G.; Kulkarni, M. V.; Venuganti, V. V. K. Dissolvable Microneedle Patch Containing Doxorubicin and Docetaxel Is Effective in 4T1 Xenografted Breast Cancer Mouse Model. Int. J. Pharm. 2019, 556, 263–275. 10.1016/j.ijpharm.2018.12.022.

(30) Janardhana, R. D.; Cabanlong, M.; Muttil, P.; Jackson, N. Barbed Microneedle Design to Enhance Penetration and Retraction Forces Using Finite Element Modelling; American Society of Mechanical Engineers Digital Collection, 2023. 10.1115/IMECE2022-95590.

(31) Zhang, Q.; Xu, C.; Lin, S.; Zhou, H.; Yao, G.; Liu, H.; Wang, L.; Pan, X.; Quan, G.; Wu, C. Synergistic Immunoreaction of Acupuncture-like Dissolving Microneedles Containing Thymopentin at Acupoints in Immune-Suppressed Rats. Acta Pharm. Sin. B 2018, 8 (3), 449–457. 10.1016/j.apsb.2017.12.006.

(32) Plotkin, S.; Robinson, J. M.; Cunningham, G.; Iqbal, R.; Larsen, S. The Complexity and Cost of Vaccine Manufacturing – An Overview. Vaccine 2017, 35 (33), 4064–4071. 10.1016/j.vaccine.2017.06.003.

(33) Kathuria, H.; Kang, K.; Cai, J.; Kang, L. Rapid Microneedle Fabrication by Heating and Photolithography. Int. J. Pharm. 2020, 575, 118992. 10.1016/j.ijpharm.2019.118992.

(34) Vassilieva, E. V.; Kalluri, H.; McAllister, D.; Taherbhai, M. T.; Esser, E. S.; Pewin, W. P.; Pulit-Penaloza, J. A.; Prausnitz, M. R.; Compans, R. W.; Skountzou, I. Improved Immunogenicity of Individual Influenza Vaccine Components Delivered with a Novel Dissolving Microneedle Patch Stable at Room Temperature. Drug Deliv. Transl. Res. 2015, 5 (4), 360–371. 10.1007/s13346-015-0228-0.

(35) Lequime, S.; Lambrechts, L. Vertical Transmission of Arboviruses in Mosquitoes: A Historical Perspective. Infect. Genet. Evol. 2014, 28, 681–690. 10.1016/j.meegid.2014.07.025.

(36) Wikan, N.; Smith, D. R. Zika Virus: History of a Newly Emerging Arbovirus. Lancet Infect. Dis. 2016, 16 (7), e119–e126. 10.1016/S1473-3099(16)30010-X.

(37) Bhatt, S.; Gething, P. W.; Brady, O. J.; Messina, J. P.; Farlow, A. W.; Moyes, C. L.; Drake, J. M.; Brownstein, J. S.; Hoen, A. G.; Sankoh, O.; Myers, M. F.; George, D. B.; Jaenisch, T.; Wint, G. R. W.; Simmons, C. P.; Scott, T. W.; Farrar, J. J.; Hay, S. I. The Global Distribution and Burden of Dengue. Nature 2013, 496 (7446), 504–507. 10.1038/nature12060.

(38) Lefteri, D. A.; Bryden, S. R.; Pingen, M.; Terry, S.; McCafferty, A.; Beswick, E. F.; Georgiev, G.; Van der Laan, M.; Mastrullo, V.; Campagnolo, P.; Waterhouse, R. M.; Varjak, M.; Merits, A.; Fragkoudis, R.; Griffin, S.; Shams, K.; Pondeville, E.; McKimmie, C. S. Mosquito Saliva Enhances Virus Infection through Sialokinin-Dependent Vascular Leakage. Proc. Natl. Acad. Sci. 2022, 119 (24), e2114309119. 10.1073/pnas.2114309119.

(39) Suh, H.; Shin, J.; Kim, Y.-C. Microneedle Patches for Vaccine Delivery. Clin. Exp. Vaccine Res. 2014, 3 (1), 42–49. 10.7774/cevr.2014.3.1.42.

(40) Sender, R.; Weiss, Y.; Navon, Y.; Milo, I.; Azulay, N.; Keren, L.; Fuchs, S.; Ben-Zvi, D.; Noor, E.; Milo, R. The Total Mass, Number, and Distribution of Immune Cells in the Human Body. Proc. Natl. Acad. Sci. U. S. A. 120 (44), e2308511120. 10.1073/pnas.2308511120.

(41) Tumban, E.; Peabody, J.; Tyler, M.; Peabody, D. S.; Chackerian, B. VLPs Displaying a Single L2 Epitope Induce Broadly Cross-Neutralizing Antibodies against Human Papillomavirus. PloS One 2012, 7 (11), e49751. 10.1371/journal.pone.0049751.

(42) Ramakanth, D.; Singh, S.; Maji, P. K.; Lee, Y. S.; Gaikwad, K. K. Advanced Packaging for Distribution and Storage of COVID-19 Vaccines: A Review. Environ. Chem. Lett. 2021, 19 (5), 3597–3608. 10.1007/s10311-021-01256-1.

(43) Tumban, E.; Muttil, P.; Escobar, C. A. A.; Peabody, J.; Wafula, D.; Peabody, D. S.; Chackerian, B. Preclinical Refinements of a Broadly Protective VLP-Based HPV Vaccine Targeting the Minor Capsid Protein, L2. Vaccine 2015, 33 (29), 3346–3353. 10.1016/j.vaccine.2015.05.016.

(44) Rouphael, N. G.; Paine, M.; Mosley, R.; Henry, S.; McAllister, D. V.; Kalluri, H.; Pewin, W.; Frew, P. M.; Yu, T.; Thornburg, N. J.; Kabbani, S.; Lai, L.; Vassilieva, E. V.; Skountzou, I.; Compans, R. W.; Mulligan, M. J.; Prausnitz, M. R.; Beck, A.; Edupuganti, S.; Heeke, S.; Kelley, C.; Nesheim, W. The Safety, Immunogenicity, and Acceptability of Inactivated Influenza Vaccine Delivered by Microneedle Patch (TIV-MNP 2015): A Randomised, Partly Blinded, Placebo-Controlled, Phase 1 Trial. The Lancet 2017, 390 (10095), 649–658. 10.1016/S0140-6736(17)30575-5.

(45) Lee, K. J.; Jeong, S. S.; Roh, D. H.; Kim, D. Y.; Choi, H.-K.; Lee, E. H. A Practical Guide to the Development of Microneedle Systems – In Clinical Trials or on the Market. Int. J. Pharm. 2020, 573, 118778. 10.1016/j.ijpharm.2019.118778.

(46) Norman, J. J.; Brown, M. R.; Raviele, N. A.; Prausnitz, M. R.; Felner, E. I. Faster Pharmacokinetics and Increased Patient Acceptance of Intradermal Insulin Delivery Using a Single Hollow Microneedle in Children and Adolescents with Type 1 Diabetes. Pediatr. Diabetes 2013, 14 (6), 459–465. 10.1111/pedi.12031.

